# Chemoinformatic-guided engineering of polyketide synthases

**DOI:** 10.1101/805671

**Authors:** Amin Zargar, Ravi Lal, Luis Valencia, Jessica Wang, Tyler William H. Backman, Pablo Cruz-Morales, Ankita Kothari, Miranda Werts, Andrew R. Wong, Constance B. Bailey, Arthur Loubat, Yuzhong Liu, Yan Chen, Veronica T. Benites, Samantha Chang, Amanda C. Hernández, Jesus F. Barajas, Mitchell G. Thompson, Carolina Barcelos, Rasha Anayah, Hector Garcia Martin, Aindrila Mukhopadhyay, Christopher J. Petzold, Edward E.K. Baidoo, Leonard Katz, Jay D. Keasling

## Abstract

Polyketide synthase (PKS) engineering is an attractive method to generate new molecules such as commodity, fine and specialty chemicals. A significant challenge in PKS design is engineering a partially reductive module to produce a saturated β-carbon through a reductive loop exchange. In this work, we sought to establish that chemoinformatics, a field traditionally used in drug discovery, could provide a viable strategy to reductive loop exchanges. We first introduced a set of donor reductive loops of diverse genetic origin and chemical substrate structures into the first extension module of the lipomycin PKS (LipPKS1). Product titers of these engineered unimodular PKSs correlated with atom pair chemical similarity between the substrate of the donor reductive loops and recipient LipPKS1, reaching a titer of 165 mg/L of short chain fatty acids produced by *Streptomyces albus* J1074 harboring these engineered PKSs. Expanding this method to larger intermediates requiring bimodular communication, we introduced reductive loops of divergent chemosimilarity into LipPKS2 and determined triketide lactone production. Collectively, we observed a statistically significant correlation between atom pair chemosimilarity and production, establishing a new chemoinformatic method that may aid in the engineering of PKSs to produce desired, unnatural products.

---

As the architecture of Type I PKSs determines the molecular product, rational reprogramming of PKS enzymes for the biosynthesis of new polyketides has been a major research thrust over the past three decades.^1–3^ Like fatty acid synthases, PKSs extend the growing chain from the ketosynthase (KS) domain with a malonyl-CoA analog loaded onto the acyl carrier protein (ACP) by the acyltransferase (AT) domain through a decarboxylative Claisen condensation reaction. Unlike fatty acid synthases, which faithfully produce saturated fatty acids, PKSs have variability in β-carbonyl reduction. After chain extension, the β-carbonyl reduction state is determined by the reductive domains within the module, namely the ketoreductase (KR), dehydratase (DH), and enoylreductase (ER), which generate the β-hydroxyl, α-β alkene, or saturated β-carbons respectively, when progressively combined. As the degree of β-carbon reduction is an important feature in molecular design, multiple studies have reported the engineering of a PKS module for various oxidation states of the β-carbon.^4–8^ However, design strategies for introduction of reductive loop exchanges (*i.e*. KR-DH-ER domains) into partially reductive modules remain elusive. In this work, we compare bioinformatic and chemoinformatic approaches to guide reductive loop (RL) exchanges and develop a new method for RL exchanges based on the chemical similarity of the RL substrate.

Chemoinformatics, an interdisciplinary field blending computational chemistry, molecular modeling and statistics, was initially developed for drug discovery through analysis of structureactivity relationships.^9^ Recently, we suggested that a chemoinformatic approach to PKS engineering could be valuable, particularly in RL exchanges due to the dependence of the KR and DH domains on substrate size.^1^ For example, a critical factor for dehydration in both standalone DH^10^ and full PKS module studies^7^ is acyl chain length, likely residing in a hydrophobic catalytic tunnel.^11,12^ Moreover, a previous study of engineered RL swaps resulted in a correlation between production and substrate size similarity of the donor RLs and the recipient module.^13^ Chemoinformatic methods such as atom pair (AP) similarity and maximum common substructure (MCS) similarity could be used to describe the substrate profiles for catalysis by these domains. AP similarity characterizes atom pairs (*e.g*. length of bond path, number of π electrons), and MCS similarity is based on identifying the largest common substructure between two molecules.^14^ While divergent in chemical characterization, both similarity methods can be translated to a Tanimoto coefficient with a range of 0 (least similar) to 1 (most similar).^14^ Based on the substrate-dependence of the reductive domains, we hypothesized that chemosimilarity between the substrates of donor and acceptor modules in RL exchanges may correlate with production levels thereby leading to engineered modules that better control the reductive state of the β carbon.

Bioinformatic studies of PKS evolution have guided engineering efforts in closely related biosynthetic gene clusters (BGCs).^15,16^ We therefore undertook a phylogenetic analysis of the reductive domain common to all RLs, the ketoreductase (KR). The KR not only reduces the β-keto group to a β-hydroxyl, but also sets the stereochemistry of the β-group and, if a branched extender is used, sets the α-carbon stereochemistry resulting in subtypes A1, A2, B1, B2 (**FigureS1A**). We generated a phylogenetic tree from all manually curated ketoreductases and ketosynthases in ClusterCAD, an online database and toolkit for Type I PKSs, totaling 72 biosynthetic gene clusters (BGCs) and 1077 modules (**FigureS1B**).^17^ As in previous investigations,^18,19^ we found that KR domains clustered by the presence of DH or DH-ER domains (**FigureS2**). We also found that the type of reductive domains do not phylogenetically cluster with the same modules KS domain or the downstream modules KS domain (**FigureS3-S4**).^18^ This suggests a link between KR evolution and product specificity, analogous to the evolution of KS domains of cis-AT^18^ and trans-AT PKS modules^20,21^ towards substrate specificity. As KRs from KR-DH-ER modules evolved distinctly from KR-only modules, we hypothesized that neither KR sequence identity or phylogenetic distance, a pairwise comparison between members of the phylogenetic tree, between the donor loops and acceptor module in RL exchanges were likely to correlate with production levels.

To compare the importance of chemical similarity and phylogenetic distance in RL exchanges, we swapped diverse RLs into the first module of the lipomycin PKS (LipPKS1) as the acceptor module. In our previous work, we introduced a heterologous thioesterase from 6-deoxyerythronolide B synthase (DEBS) into the C-terminus of LipPKS1; the resulting truncated PKS produced a β-hydroxy acid.^22^ In this work, our initial experimental design was based on introducing full RLs using conserved residues into this module as exchange sites, generating a saturated acid (**Scheme1**).^7^ We selected N-terminal junctions (“A” and “B”) located immediately after the post-AT linker, which is important for KS-AT domain architecture,^23^ and the C-terminal junction (“C”) directly before the ACP domain (see **TableS1** for sequences) based on our previous work in the first module of borreledin.^7^

**Scheme 1.**
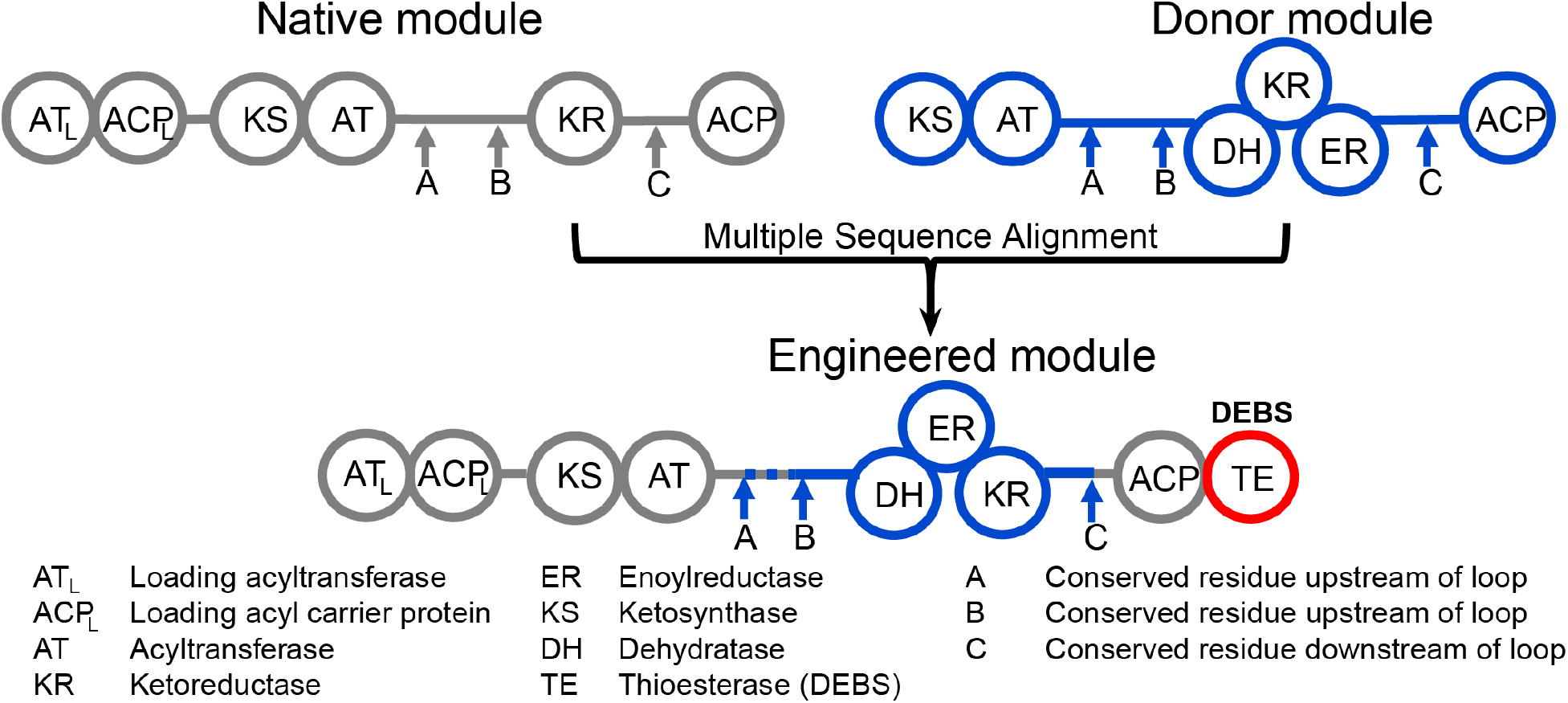
Experimental design of RL swaps. Conserved residues are identified through multiple sequence alignment surrounding the reductive domains (“A”, “B” and “C”). Donor RLs are inserted into the native lipomycin module 1, and the attached DEBS thioesterase hydrolyzes the product.

To evaluate the effects of genetic and chemical similarity, we identified five donor RLs (IdmO, indanomycin, *S. antibioticus;* SpnB, spinosyn, *S. spinosa;* AurB, aureothin, *S. aureofaciens;* NanA2, nanchangamycin, *S. nanchangensis;* final products in **FigureS5**) to swap into LipPKS1. A pairwise comparison of phylogenetic distance and amino acid sequence identity determined that the KRs of IdmO, AurB, and SpnB are the most similar to LipPKS1 (**Figure1A**). A similar trend holds in the analysis of each of these donor modules KS domain and downstream KS domain compared to the KS of LipPKS1 (**FigureS6**). In contrast, the NanA2 substrate is the most chemically similar to LipPKS1, followed by SpnB, based on AP and MCS similarity (**Figure1B**). With the introduction of a RL swap, the chimeric enzymes should produce 2,4-dimethyl pentanoic acid. As *in vitro* PKS studies have shown divergence from *in vivo* results^24,25^ due to underestimation of factors including limiting substrate, crowding, and solubility,^26^ we cloned eight chimeric modules, along with a control expressing red fluorescent protein (RFP), into an *E. coli -Streptomyces albus* shuttle vector and conjugated it into *S. albus* J1074 (**TableS1**).^27^ Following ten-day production runs in a rich medium in biological triplicate, cultures of *S. albus* harboring each of the constructs were harvested and the supernatants were analyzed for product levels (**Supplemental Methods**).

**Figure 1.**
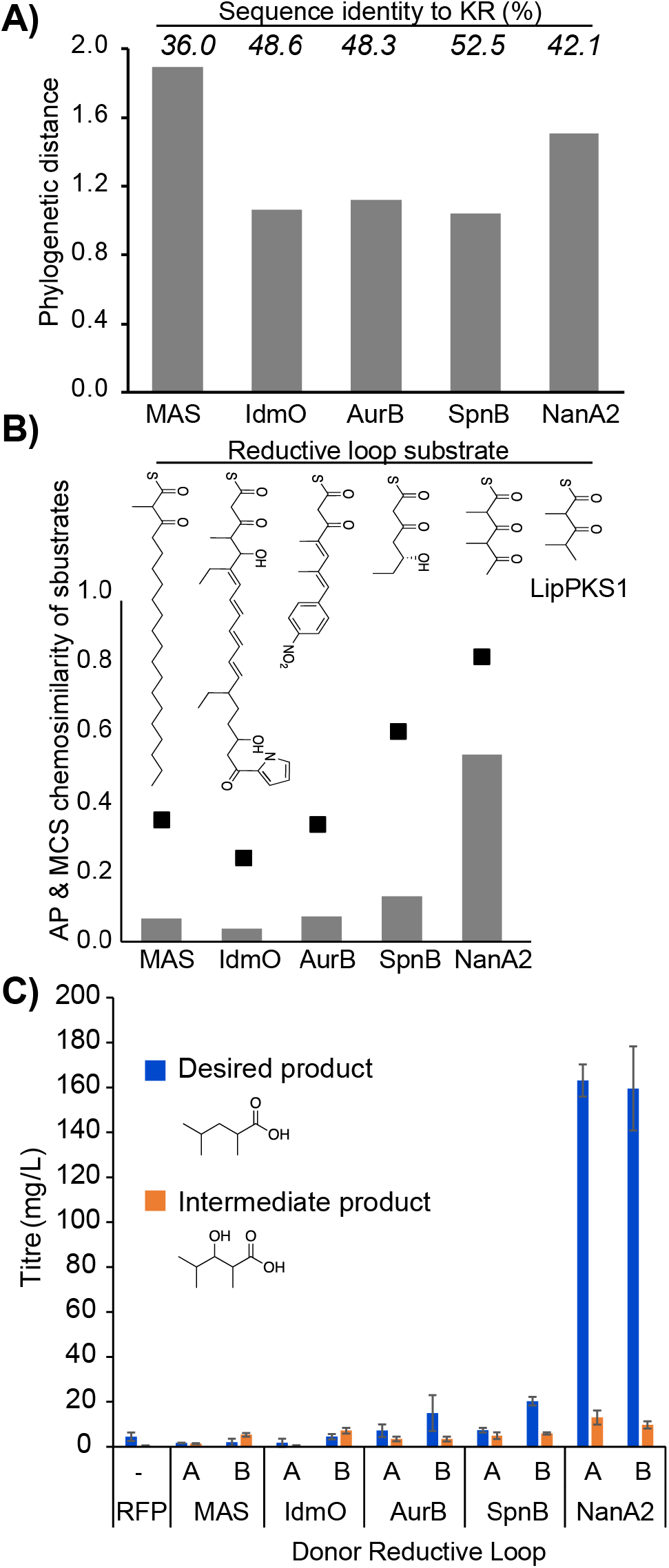
Phylogenetic and chemical similarity effects on reductive loop exchanges. **A)** Phylogenetic distance of the native LipPKS1 KR domain to each donor KR. The value above each bar denotes KR sequence identity comparison. **B)** AP (bar) and MCS (dots) chemical similarity between the native LipPKS1 KR domain and each of the donor KR domains. Chemical structures display native KR substrate in each module **C)** Polyketide production of engineered PKSs at both junction “A” and junction “B” in biological triplicate (error bars denote standard deviation).

Consistent with our hypothesis, we found a perfect correlation between titers of the desired product and the AP and MCS chemosimilarities of the donor and LipPKS1 module substrates (Rs=1.00 and p=0.00) (**Figure1C**). On the other hand, no significant correlation between product titer and phylogenetic distance or sequence similarity of the KR domain (Rs=0.04, p=0.60) was found. Based on our bioinformatic analysis, this was not surprising as the lipomycin KR is an A2-type, evolving separately from a KR with a full RL. This trend held with either junction A or B, although generally junction B chimeras resulted in higher product titers, consistent with a previous study of RL exchanges, as the extra residues in junction A are distal to the ACP docking interface and active site.^7^ We found that substituting the donor loop most chemically similar to LipPKS1, NanA2, resulted in the highest titers of the desired product, 2,4-dimethyl pentanoic acid, reaching 165 mg/L (Supplemental Methods). Low titers of the intermediate 2,4-dimethyl-3-hydroxypentanoic acid were produced, which we hypothesize is due to a comparatively lower rate of turnover at the energetically intensive DH domain,^28^ resulting in premature cleavage of the stalled product by non-enzymatic or TE-mediated hydrolysis. As in our previous study of *in vitro* production of adipic acid, we did not detect alkene or keto acid stalled products^7^. This is not surprising as non-functional KRs produce short chain β-keto acids that spontaneously decarboxylate to form ketones, which we did not observe, and ERs have been generally shown to rapidly reduce *trans* double bonds.^28^

Based on these results, we took a chemoinformatic approach to further test our hypothesis that chemosimilarity of RL substrates is a critical factor in PKS engineering. We searched the ClusterCAD^17^ database for PKS modules with full RLs and substrates of high chemical similarity to that of the KR of LipPKS1. The closest matches identified were the second PKS module from laidlomycin and monensin, which have the same substrate as NanA2 (**Figure2A**). As junction B resulted in levels of production superior to junction A, we cloned the RLs of LaidS2 and MonA2 into junction B of lipomycin. Like NanA2, LaidS2 loops produced much higher titers of the fully desired product, while MonA2 produced at similar levels to SpnB and AurB (**Figure2B**). As protein levels may influence product titers, we determined the quantitative levels of all LipPKS1 constructs using targeted proteomics at the conclusion of the production run and observed no correlation between PKS protein levels and product titers (Rs = -.15 and p=.77) (**FigureS7**). We observed reduced protein levels in the MonA2 swap, which could partially explain the lower levels of production in the MonA2 swap compared to LaidS2 and NanA2. We should note however that targeted proteomics of three peptide peaks across the PKS does not eliminate the possibility of proteolytic degradation nor variability in protein quality. We determined that AP Tanimoto and MCS chemosimilarity had equivalent Spearman rank correlation to product titers (Rs of 0.82, p = 0.05).

**Figure 2.** A chemoinformatic approach to reductive loop exchanges. **A)** ClusterCad search revealed the closest substrates to LipPKS1 containing a full RL **B)** Production levels of junction “B” RL exchanges ordered from most similar KR substrate to LipPKS1 (MonA2, LaidS2 and NanA2) to progressively less similarity (IdmO, AurB, SpnB) in biological triplicate (error bars denote standard deviation).

We sought to better demonstrate the utility of this approach through RL exchanges where AP and MCS chemosimilarity diverge. Moreover, we wanted to test this method with a module located in the center of an assembly line, which requires docking domain interactions and a larger substrate. We therefore performed RL swaps on the second module of lipomycin, LipPKS2 (**Figure3A**), to generate triketide lactones. We selected donor loops from SpnB and NanA2, as NanA2 has higher AP chemosimilarity while SpnB has higher MCS chemosimilarity (**Figure3B**). We found higher product levels using NanA2 than SpnB (**Figure3C-D**), correlating better with AP chemical similarity than with MCS chemical similarity, and importantly, as in our single-module swaps, KR phylogenetic similarity and sequence identity did not correlate with product titers. We should note that proteomics on each PKS of these bimodular systems was not performed to rule out the effect of variable protein levels. As AP chemosimilarity more heavily weights substructures, this metric produces higher levels of similarity between NanA2 and LipPKS2 because both select methylmalonyl-CoA in the first two modules. In contrast, as MCS chemosimilarity simply considers the largest common substructure, malonyl-selecting SpnB is considered more similar, ignoring that commonality at the growing chain by methyl groups may be more influential than more distal regions of the substrate. While extension of this phenomenon to account for variances in chemical similarity metrics (*e.g*. AP, MCS) requires further study, we hypothesize that chemosimilarity metrics that best match PKS enzymatic processing may prove most successful. Overall, in our reductive loop exchanges we determined a Spearman correlation between AP Tanimoto chemosimilarity and product titer to have an Rs of 0.88 and a p-value of 0.004 (**Supplemental Methods**).

**Figure 3.**
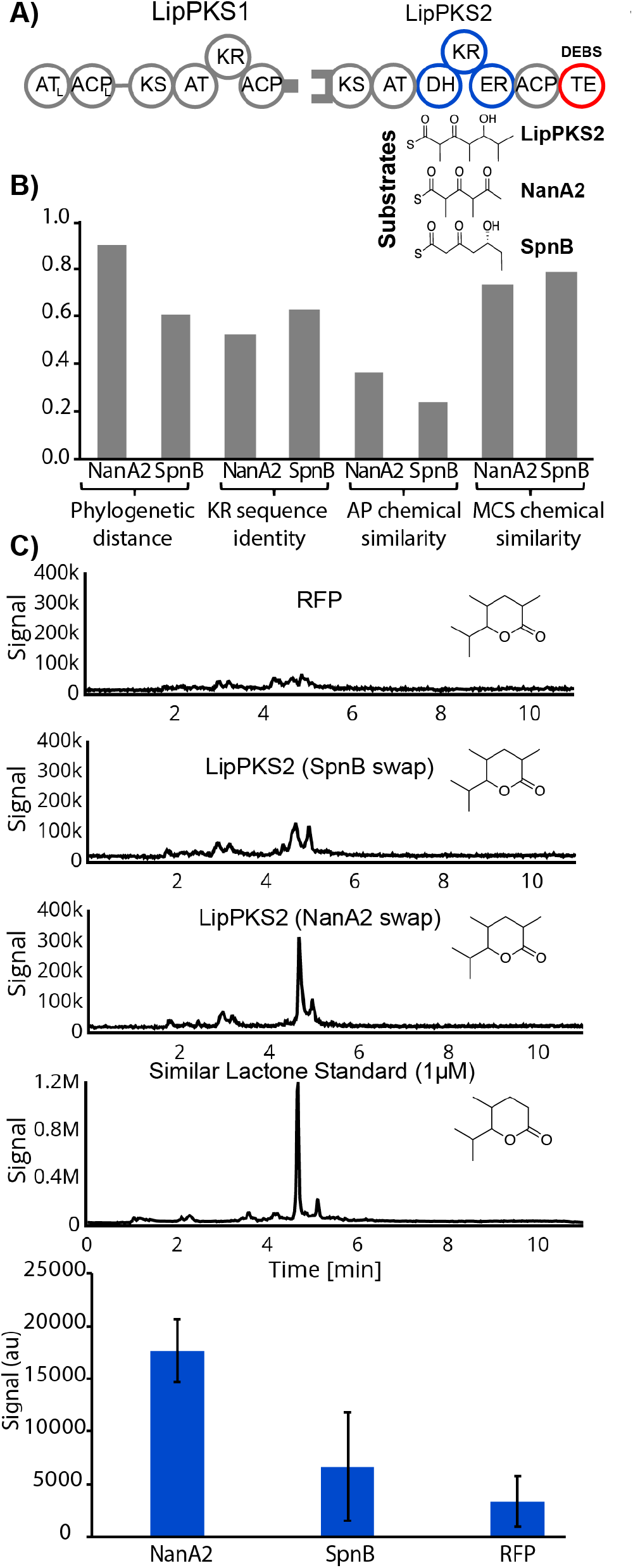
Bimodular reductive loop exchange **A)** Schematic of reductive loop exchanges in LipPKS2 with substrates **B)** Phylogenetic distance, KR sequence identity, AP chemosimilarity and MCS similarity between reductive loop donors and LipPKS2 **C)** Chromatograms of RFP, LipPKS2 with donor loops SpnB and NanA2, and a structurally similar standard spiked into RFP cultures **D)** Production levels of desired triketide lactone in biological triplicate (error bars denote standard deviation).

Highlighting previous literature regarding the importance of substrate size in reductive domains, we hypothesized that the field of chemoinformatics, traditionally used to study structure-activity relationships in drug discovery, could be applied to PKS engineering. Using different RLs of varying phylogenetic and chemical similarity, we determined that chemosimilarity between donor KRs and recipient KRs correlated with production, in contrast to phylogenetic distance and sequence similarity. Extending our method into multi-modular systems that use larger substrates and communication domains, we performed RL swaps in LipPKS2 and found that AP chemosimilarity correlates with production. While our approach did not find a correlation between genetic similarity and production in RL swaps, it has been shown that within highly similar BGCs, the downstream KS clades with the extent of upstream reduction^16^. In this study, the donor modules do not share close homology with the lipomycin recipient module, and we believe it likely that donor loops with high chemosimilarity located within a BGC would prove more compatible than chemosimilarity alone. Overall, our results determined statistical significance in the correlation between production and the chemosimilarity of the substrate between the donor and recipient modules. More generally, chemoinformatics may provide guideposts for other engineering goals (*e.g*. KR domain subtype swaps to switch stereochemistry). With our incomplete understanding of PKS processing, design principles may accelerate the combinatorial approach currently used for *de novo* biosynthesis and help provide a framework to more rapidly produce valuable biochemicals.

## Acknowledgements

This work was funded by the DOE Joint BioEnergy Institute (http://www.jbei.org) supported by the U.S. Department of Energy, Office of Science, Office of Biological and Environmental Research between Lawrence Berkeley National Laboratory and the U.S. Department of Energy, the Agile Biofoundry sponsored by the U.S. DOE Office of Energy Efficiency and Renewable Energy, Bioenergy Technologies and Vehicle Technologies Offices, under Contract DEAC02-05CH11231 between DOE and Lawrence Berkeley National Laboratory, and the National Institute of Health Awards F32GM125179, F32GM125166. H.G.M. was also supported by the Basque Government through the BERC 2018-2021 program and by Spanish Ministry of Economy and Competitiveness MINECO: BCAM Severo Ochoa excellence accreditation SEV-2017-0718.

## Competing Financial Interest

J.D.K. has a financial interest in Amyris, Lygos, Demetrix, Napigen, Maple Bio, Berkeley Brewing Sciences, Ansa Biotech and Apertor Labs.

## Supporting Information

Supplementary tables and figures, and detailed experimental procedures

## References

(1) Barajas, J. F.; Blake-Hedges, J. M.; Bailey, C. B.; Curran, S.; Keasling, J. D. Synergy between protein and host level engineering. Synthetic and Systems Biotechnology 2017, 2, 147–166.

(2) Khosla, C.; Herschlag, D.; Cane, D. E.; Walsh, C. T. Assembly line polyketide synthases: mechanistic insights and unsolved problems. Biochemistry 2014, 53, 2875–2883.

(3) Yuzawa, S.; Zargar, A.; Pang, B.; Katz, L.; Keasling, J. D. Commodity chemicals from engineered modular type I polyketide synthases. 2018, 608, 393–415.

(4) Reid, R.; Piagentini, M.; Rodriguez, E.; Ashley, G.; Viswanathan, N.; Carney, J.; Santi, D. V.; Hutchinson, C. R.; McDaniel, R. A model of structure and catalysis for ketoreductase domains in modular polyketide synthases. Biochemistry 2003, 42, 72–79.

(5) Keatinge-Clay, A. Crystal structure of the erythromycin polyketide synthase dehydratase. J. Mol. Biol. 2008, 384, 941–953.

(6) Kellenberger, L.; Galloway, I. S.; Sauter, G.; Böhm, G.; Hanefeld, U.; Cortés, J.; Staunton, J.; Leadlay, P. F. A polylinker approach to reductive loop swaps in modular polyketide synthases. Chembiochem 2008, 9, 2740–2749.

(7) Hagen, A.; Poust, S.; Rond, T. de; Fortman, J. L.; Katz, L.; Petzold, C. J.; Keasling, J. D. Engineering a polyketide synthase for in vitro production of adipic acid. ACS Synth. Biol. 2016, 5, 21–27.

(8) Gaisser, S.; Kellenberger, L.; Kaja, A. L.; Weston, A. J.; Lill, R. E.; Wirtz, G.; Kendrew, S. G.; Low, L.; Sheridan, R. M.; Wilkinson, B.; Galloway, I. S.; Stutzman-Engwall, K.; McArthur, H. A.; Staunton, J.; Leadlay, P. F. Direct production of ivermectin-like drugs after domain exchange in the avermectin polyketide synthase of Streptomyces avermitilis ATCC31272. Org. Biomol. Chem. 2003, 1, 2840–2847.

(9) Maldonado, A. G.; Doucet, J. P.; Petitjean, M.; Fan, B.-T. Molecular similarity and diversity in chemoinformatics: from theory to applications. Mol Divers 2006, 10, 39–79.

(10) Faille, A.; Gavalda, S.; Slama, N.; Lherbet, C.; Maveyraud, L.; Guillet, V.; Laval, F.; Quémard, A.; Mourey, L.; Pedelacq, J.-D. Insights into Substrate Modification by Dehydratases from Type I Polyketide Synthases. J. Mol. Biol. 2017, 429, 1554–1569.

(11) Herbst, D. A.; Jakob, R. P.; Zähringer, F.; Maier, T. Mycocerosic acid synthase exemplifies the architecture of reducing polyketide synthases. Nature 2016, 531, 533–537.

(12) Barajas, J. F.; McAndrew, R. P.; Thompson, M. G.; Backman, T. W. H.; Pang, B.; de Rond, T.; Pereira, J. H.; Benites, V. T.; Martín, H. G.; Baidoo, E. E. K.; Hillson, N. J.; Adams, P. D.; Keasling, J. D. Structural insights into dehydratase substrate selection for the borrelidin and fluvirucin polyketide synthases. J. Ind. Microbiol. Biotechnol. 2019, 46, 1225–1235.

(13) McDaniel, R.; Thamchaipenet, A.; Gustafsson, C.; Fu, H.; Betlach, M.; Ashley, G. Multiple genetic modifications of the erythromycin polyketide synthase to produce a library of novel “unnatural” natural products. Proc. Natl. Acad. Sci. USA 1999, 96, 1846–1851.

(14) Chen, X.; Reynolds, C. H. Performance of similarity measures in 2D fragment-based similarity searching: comparison of structural descriptors and similarity coefficients. J Chem Inf Comput Sci 2002, 42, 1407–1414.

(15) Peng, H.; Ishida, K.; Sugimoto, Y.; Jenke-Kodama, H.; Hertweck, C. Emulating evolutionary processes to morph aureothin-type modular polyketide synthases and associated oxygenases. Nat. Commun. 2019, 10, 3918.

(16) Awakawa, T.; Fujioka, T.; Zhang, L.; Hoshino, S.; Hu, Z.; Hashimoto, J.; Kozone, I.; Ikeda, H.; Shin-Ya, K.; Liu, W.; Abe, I. Reprogramming of the antimycin NRPS-PKS assembly lines inspired by gene evolution. Nat. Commun. 2018, 9, 3534.

(17) Eng, C. H.; Backman, T. W. H.; Bailey, C. B.; Magnan, C.; García Martín, H.; Katz, L.; Baldi, P.; Keasling, J. D. ClusterCAD: a computational platform for type I modular polyketide synthase design. Nucleic Acids Res. 2018, 46, D509–D515.

(18) Zhang, L.; Hashimoto, T.; Qin, B.; Hashimoto, J.; Kozone, I.; Kawahara, T.; Okada, M.; Awakawa, T.; Ito, T.; Asakawa, Y.; Ueki, M.; Takahashi, S.; Osada, H.; Wakimoto, T.; Ikeda, H.; Shin-Ya, K.; Abe, I. Characterization of Giant Modular PKSs Provides Insight into Genetic Mechanism for Structural Diversification of Aminopolyol Polyketides. Angew. Chem. Int. Ed. Engl. 2017, 56, 1740–1745.

(19) Jenke-Kodama, H.; Börner, T.; Dittmann, E. Natural biocombinatorics in the polyketide synthase genes of the actinobacterium Streptomyces avermitilis. PLoS Comput. Biol. 2006, 2, e132.

(20) Nguyen, T.; Ishida, K.; Jenke-Kodama, H.; Dittmann, E.; Gurgui, C.; Hochmuth, T.; Taudien, S.; Platzer, M.; Hertweck, C.; Piel, Exploiting the mosaic structure of trans-acyltransferase polyketide synthases for natural product discovery and pathway dissection. J. Nat. Biotechnol. 2008, 26, 225–233.

(21) Vander Wood, D. A.; Keatinge-Clay, A. T. The modules of trans-acyltransferase assembly lines redefined with a central acyl carrier protein. Proteins 2018, 86, 664–675.

(22) Yuzawa, S.; Eng, C. H.; Katz, L.; Keasling, J. D. Broad substrate specificity of the loading didomain of the lipomycin polyketide synthase. Biochemistry 2013, 52, 3791–3793.

(23) Tang, Y.; Kim, C.-Y.; Mathews, I. I.; Cane, D. E.; Khosla, C. The 2.7-Angstrom crystal structure of a 194-kDa homodimeric fragment of the 6-deoxyerythronolide B synthase. Proc. Natl. Acad. Sci. USA 2006, 103, 11124–11129.

(24) Khosla, C.; Tang, Y.; Chen, A. Y.; Schnarr, N. A.; Cane, D. E. Structure and mechanism of the 6-deoxyerythronolide B synthase. Annu. Rev. Biochem. 2007, 76, 195–221.

(25) Yan, J.; Hazzard, C.; Bonnett, S. A.; Reynolds, K. A. Functional modular dissection of DEBS1-TE changes triketide lactone ratios and provides insight into Acyl group loading, hydrolysis, and ACP transfer. Biochemistry 2012, 51, 9333–9341.

(26) Zotter, A.; Bäuerle, F.; Dey, D.; Kiss, V.; Schreiber, G. Quantifying enzyme activity in living cells. J. Biol. Chem. 2017, 292, 15838–15848.

(27) Phelan, R. M.; Sachs, D.; Petkiewicz, S. J.; Barajas, J. F.; Blake-Hedges, J. M.; Thompson, M. G.; Reider Apel, A.; Rasor, B. J.; Katz, L.; Keasling, Development of Next Generation Synthetic Biology Tools for Use in Streptomyces venezuelae. J. D. ACS Synth. Biol. 2017, 6, 159–166.

(28) Weber, A. L. Origin of fatty acid synthesis: thermodynamics and kinetics of reaction pathways. J. Mol. Evol. 1991, 32, 93–100.

