## Supplementary Info for "Chemoinformatic-guided engineering of polyketide synthases"

### Supporting Information

| **Strains** | **JBEI Number** | **Source** | **Notes** |
|---|---|---|---|
| *E. coli* ET12567 | | ATCC BAA-525 | |
| *S. venezuelae* | | ATCC 10712 | |
| **Plasmids** | | | |
| Lip1 JuncA IdmO | JBx_078437 | This study | Chimeric Lip1 with reductive loops swap from IdmO at junction A with DEBS thioesterase |
| Lip1 JuncB IdmO | JBx_078438 | This study | Chimeric Lip1 with reductive loops swap from IdmO at junction B with DEBS thioesterase |
| Lip1 JuncA SpnB | JBx_078439 | This study | Chimeric Lip1 with reductive loops swap from SpnB at junction A with DEBS thioesterase |
| Lip1 JuncB SpnB | JBx_081781 | This study | Chimeric Lip1 with reductive loops swap from SpnB at junction B with DEBS thioesterase |
| Lip1 JuncB AurB | JBx_082096 | This study | Chimeric Lip1 with reductive loops swap from AurB at junction B with DEBS thioesterase |
| Lip1 JuncA AurB | JBx_082101 | This study | Chimeric Lip1 with reductive loops swap from AurB at junction A with DEBS thioesterase |
| Lip1 JuncB NanA2 | JBx_082097 | This study | Chimeric Lip1 with reductive loops swap from NanA2 at junction B with DEBS thioesterase |
| Lip1 JuncA NanA2 | JBx_081782 | This study | Chimeric Lip1 with reductive loops swap from NanA2 at junction A with DEBS thioesterase |
| Lip1 JuncB MAS | JBx_082098 | This study | Chimeric Lip1 with reductive loops swap from MAS at junction B with DEBS thioesterase |
| Lip1 JuncA MAS | JBx_081709 | This study | Chimeric Lip1 with reductive loops swap from MAS at junction A with DEBS thioesterase |

| | | | |
|---|---|---|---|
| Lip1 JuncA MonA2 | JBx_083535 | This study | Chimeric Lip1 with reductive loops swap from MonA2 at junction A with DEBS thioesterase |
| Lip1 JuncA LaidS2 | JBx_084029 | This study | Chimeric Lip1 with reductive loops swap from LaidS2 at junction A with DEBS thioesterase |
| Lip1 Native | JBx_082455 | This study | Native Lip1 module |
| Lip2 JuncA SpnB | JBx_083532 | This study | Chimeric Lip2 with reductive loops swap from SpnB at junction A with DEBS thioesterase |
| Lip2 JuncA NanA2 | JBx_083533 | This study | Chimeric Lip2 with reductive loops swap from NanA2 at junction A with DEBS thioesterase |

**Table S1.** All strains and plasmids used in this study. All sequences are publicly available and can be physically requested at https://public-registry.jbei.org/folders/504.

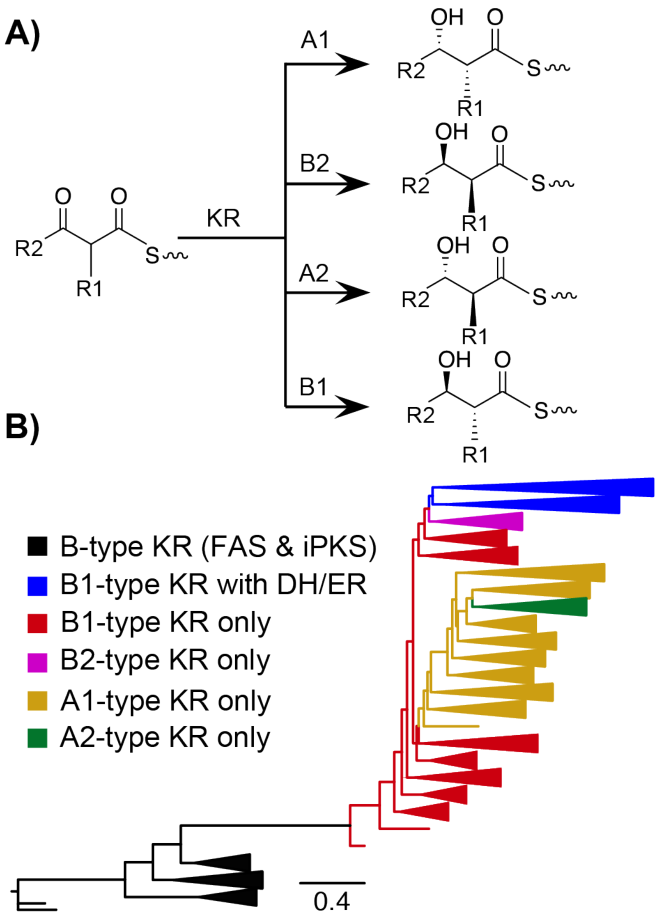


**Figure S1.** Bioinformatic analysis of reductive loop exchanges. **A)** KR subtypes determine the stereochemistry of the β-hydroxyl and α-carbon **B)** Phylogenetic tree of the ketoreductase (KR) domain of all manually curated KRs in ClusterCAD determined by ModelFinder in IQ-Tree. This evolutionary reconstruction revealed that KR-only (reductive loops with only a KR domain) B1 subtypes split from a common ancestor of fatty acid synthases and iterative PKSs. As in previous investigations, we found that KR-only B1 subtypes later resulted in the addition of DH and DH/ER domains,[18] likely through recombination.[19] We extend this finding to note that the KR-only B1 subtype branch diverged to produce the other KR-only subtypes (*i.e.* A1, A2 and B2).

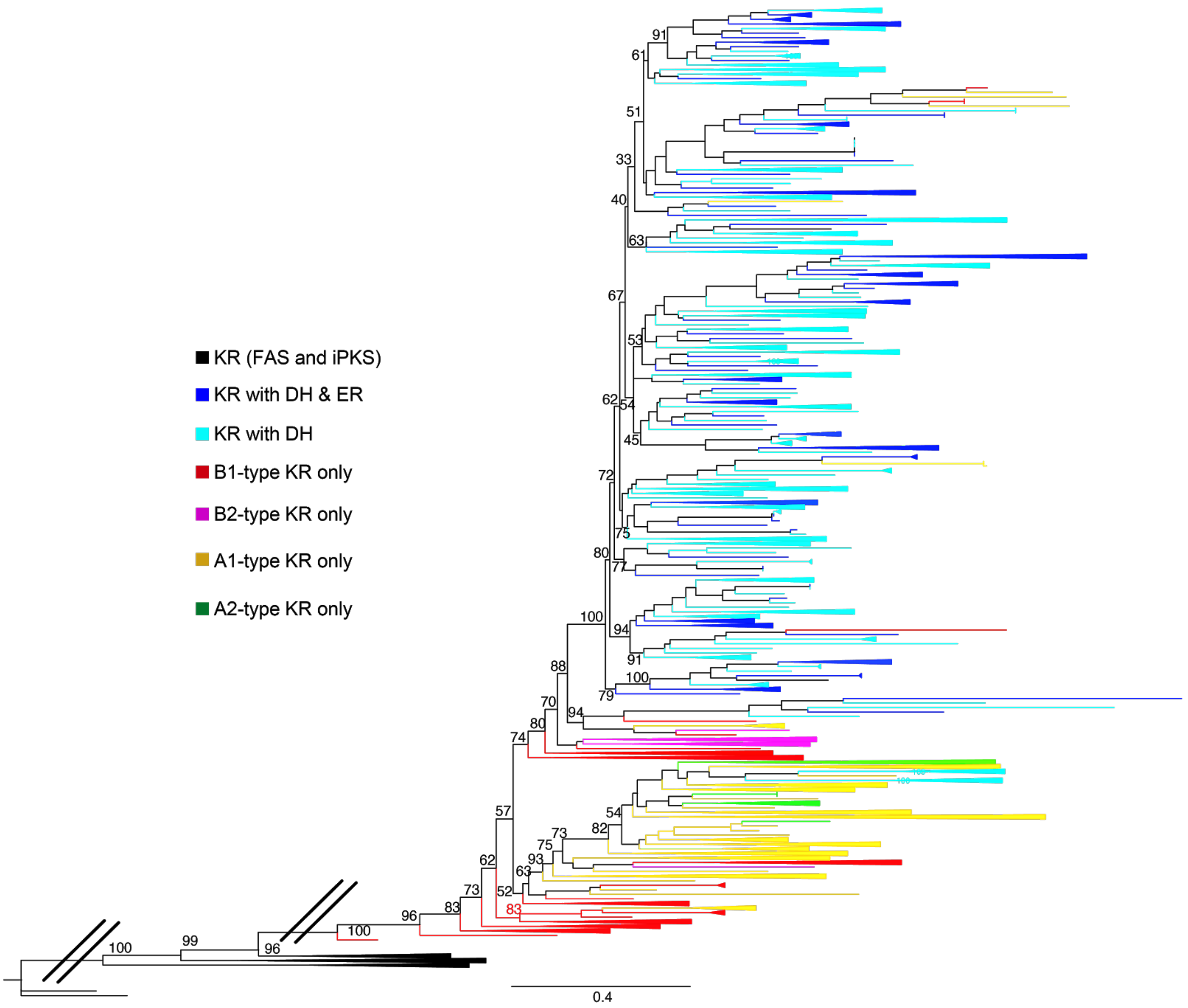


**Figure S2.** Phylogenetic tree of the ketoreductase (KR) domain. The KR domain of all modules in ClusterCad that contained a functional and annotated KR domain were manually curated were extracted. A phylogenetic tree of these classes of KR domains were built through ModelFinder in IQ-Tree. Each branch represents a KR domain and the numerical values at each branch nodes represent the branch support as a percentage value.

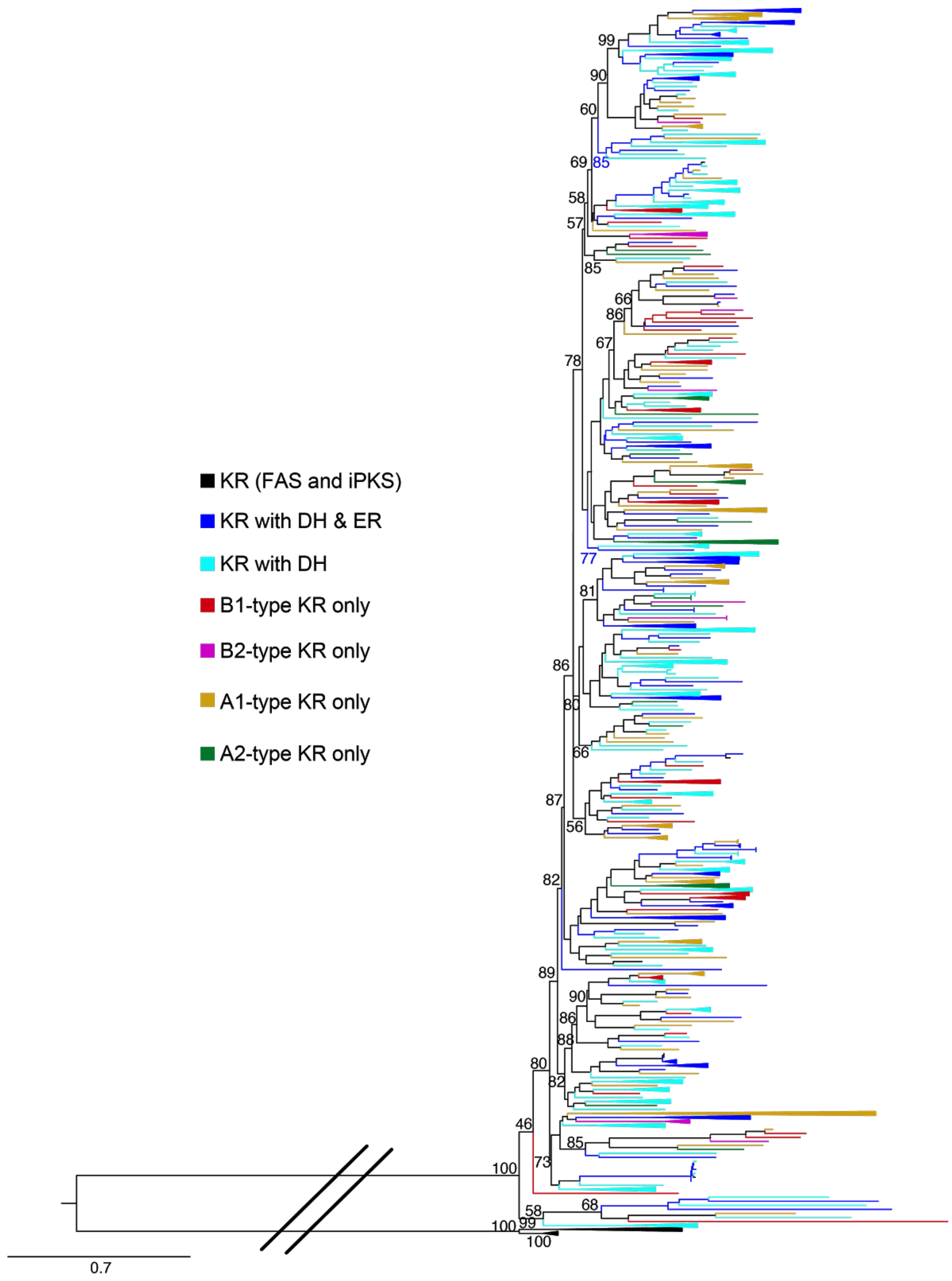


**Figure S3.** Phylogenetic tree of the ketosynthase (KS) domain. The KS domain of all modules in ClusterCad that contain a functional and annotated KR domain were manually curated were extracted. A phylogenetic tree of these KS domains from each class of KR domains were built through ModelFinder in IQ-Tree. Each branch represents a KS domain in ClusterCad and the numerical values at each branch nodes represent the branch support.

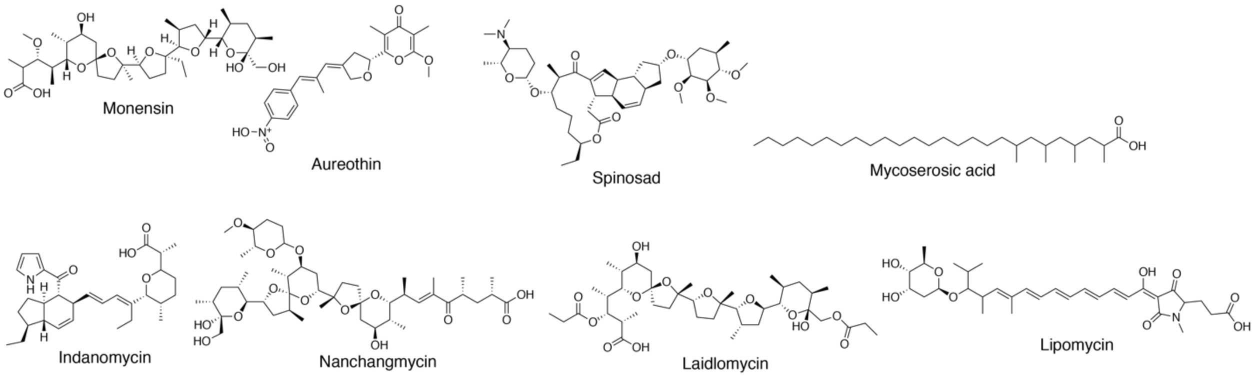


**Figure S4.** Structures of the final products of the recipient PKS (lipomycin) and the PKSs harboring the donor loops.

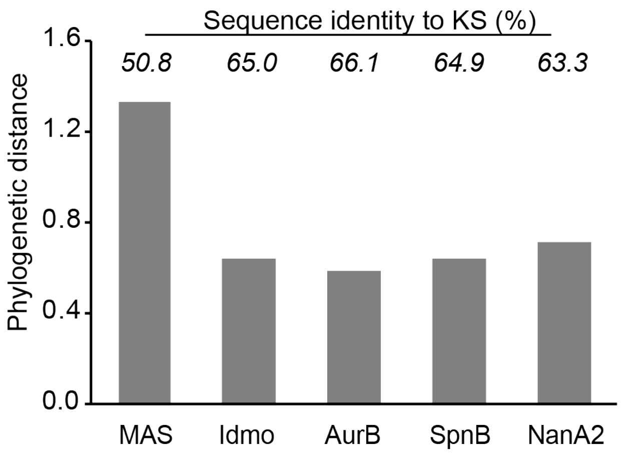


**Figure S5.** Phylogenetic similarity of the native Lip1 KS domain to each donor KS, normalized to the most similar and least similar KS domain in ClusterCad. The value above each bar denotes the sequence identity percentage.

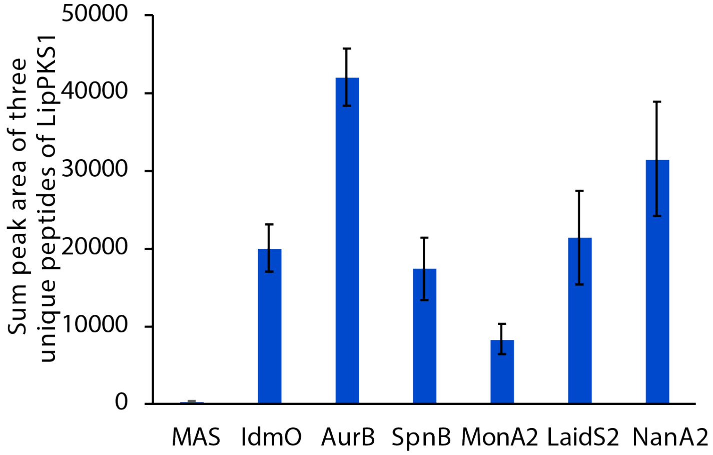


**Figure S6.** Proteomics of LipPKS1 reductive loop swaps at junction A. The cells were harvested at the end of production (day 10). Three tryptic peptides that are shared in all engineered LipPKS1 were quantified using described targeted MS method. The sum peak area of these peptides was used as total protein peak area to relatively quantify across samples expressing different LipPKS1. The mean value and standard deviation of the protein peak area in three biological replicates (N = 3) were plotted in the bar graph.

**Supplemental Methods**

Cloning of engineered LipPKS1 reductive loop modules

The phiC31 *Streptomyces* integrase vectors were used as described by Phelan *et al* to integrate the LipPKS1 reductive loop swap modules[27]. The native lipomycin module 1 with a fused DEBS thioesterase was used from Yuzawa *et al*[30]. Reductive loop sequences of IdmO, AurB, NanA2, and SpnB sequences were codon optimized for *E. coli* and amplified from Hagen *et al*[7]. Reductive loop sequences from MAS, MonA2, and LaidS2 were codon optimized for *E. coli* and synthesized by Gen9 (since acquired by Ginkgo Bioworks). Cloning was performed through Golden Gate assembly. All clusters were expressed under the GapDH(El) promoter from *Eggerthella lenta.* Junction sites for reductive loop sites were determined by those reported by Hagen *et al* through multiple sequence alignment with Muscle[31]. The plasmids along with their associated information have been deposited in the public version of JBEI registry (http://public-registry.jbei.org) and are physically available from the authors upon request https://public-registry.jbei.org/folders/504.

Cloning of native LipPKS1 and LipPKS2 reductive loop modules

The phiC31 integrase vectors were used to integrate the native LipPKS1 module. The cloning of the native docking domain to replace the DEBS thioesterase was performed through Golden Gate assembly. The VWB *Streptomyces* integrase vectors were used as described by Phelan *et al* [1] to integrate the LipPKS2 reductive loop swap modules. The native LipPKS2 was codon optimized for *E. coli* and the native sequences synthesized with an attached DEBS thioesterase. Junction A reductive loop sites were determined through Muscle as in LipPKS1, and SpnB and NanA2 reductive loops were cloned through Golden Gate assembly. The plasmids along with their associated information have been deposited in the public version of JBEI registry (http://public-registry.jbei.org) and are physically available from the authors upon request https://public-registry.jbei.org/folders/504.

Conjugation of phiC31 integrase LipPKS1 constructs into *Streptomyces albus*

*E. coli* ET12567/pUZ8002 was transformed with LipPKS1 plasmids and selected for on LB agar containing kanamycin (25 µg/mL), chloramphenicol (15 µg/mL), and apramycin (50 µg/mL). A single colony was used to inoculate a 5 mL of LB containing kanamycin (25 µg/mL), chloramphenicol (15 µg/mL), and apramycin (50 µg/mL) at 37°C. The overnight culture was used to seed 10 mL of LB containing the same antibiotics, which was grown at 37°C to an OD600 of 0.4-0.6. The *E. coli* cells were pelleted by centrifugation, washed twice with LB, and resuspended in 500 µL of LB. Fresh *S. albus* J1074 spores were collected from a mannitol soy agar plate with 5 mL of 2xYT and incubated at 50°C for 10 min. The spores (500 µL) and the *E. coli* cells (500 µL) were mixed, spread onto mannitol soy agar, and incubated at 30°C for 16 hours. After 1 mL addition of nalidixic acid (20 µg/mL) and apramycin (40 µg/mL) was added and allowed to dry, the plate was further incubated for 3-4 days at 30°C. A single colony was used to inoculate into TSB containing nalidixic acid (25 µg/mL) and apramycin (25 µg/mL). After 3-4 days, a 1 mL aliquot was taken for genomic isolation through the Maxwell kit (Promega, Cat# AS1490, Madison WI). Successful integration was verified through qPCR. The remainder of the culture was spread onto a MS plate and incubated at 30°C for 2-3 days. The spores were collected from the plate with 3-4 mL of water and mixed with glycerol to prepare 25% glycerol stock. The glycerol stock was stored at -80°C for long-term storage.

Conjugation of VWB integrase LipPKS2 constructs into *Streptomyces albus*

*E. coli* ET12567/pUZ8002 was transformed with LipPKS2 plasmids and selected for on LB agar containing kanamycin (25 µg/mL), chloramphenicol (15 µg/mL), and spectinomycin (100 µg/mL). A single colony was used to inoculate a 5 mL of LB containing kanamycin (25 µg/mL), chloramphenicol (15 µg/mL), and apramycin (100 µg/mL) at 37°C. The overnight culture was used to seed 10 mL of LB containing the same antibiotics, which was grown at 37°C to an OD600 of 0.4-0.6. The *E. coli* cells were pelleted by centrifugation, washed twice with LB, and resuspended in 500 µL of LB. *S. albus* J1074 spores with an integrated LipPKS1 were collected from a mannitol soy agar plate with 5 mL of 2xYT and incubated at 50°C for 10 min. The spores (500 µL) and the *E. coli* cells (500 µL) were mixed, spread onto mannitol soy agar, and incubated at 30°C for 16 hours. After a 1-mL addition of nalidixic acid (20 µg/mL), apramycin (40 µg/mL), and spectinomycin (400 µg/mL) was added and allowed to dry, the plate was further incubated for 3-4 days at 30°C. A single colony was used to inoculate into TSB containing nalidixic acid (25 µg/mL), apramycin (25 µg/mL) and spectinomycin (100 µg/mL). After 3-4 days, a 1-mL aliquot was taken for genomic isolation through the Maxwell kit (Promega, Cat# AS1490, Madison WI). Successful integration was verified through qPCR. The remainder of the culture was spread onto a MS plate and incubated at 30ºC for 2-3 days. The spores were collected from the plate with 3-4 mL of water and mixed with glycerol to prepare 25% glycerol stock. The glycerol stock was stored at -80°C for long-term storage.

*S. albus* production runs

Engineered *S. albus* spores were grown in 12 mL of TSB medium containing nalidixic acid (50 µg/mL) and apramycin (50 µg/mL) for 4-5 days at 30°C for single module LipPKS1 studies. Bimodular studies also included spectinomycin (200 µg/mL). 3 mL of the overnight culture was used to seed 30 mL of 10% media 042 and 90% plant hydrolysate[5], supplemented with 2.4 grams/liter of valine and nalidixic acid (50 µg/mL), which was grown for 10 days at 30°C. For bimodular production runs, an overlay of 4 mL of dodecane was added to retain the triketide lactone.

Sample preparation for single module production of short chain acids

To detect β-keto, β-hydroxy, a-β alkene, and saturated acids, 1 mL of each sample was centrifuged at 5000xg for 10 minutes and 200 µL of the supernatant was removed. The supernatant was mixed with 200 µl of 100 µM hexanoic acid dissolved in methanol and filtered using Amicon Ultra Centrifugal filters, 3 kDa Ultracel, 0.5 mL device (Millipore). β-hydroxy (3-hydroxy- 2,4-dimethylpentanoic acid) and saturated acids (2,4-dimethylpentanoic acid) were synthesized by Enamine (Cincinnati, USA) to greater than 95% purity.

Sample preparation for bimodular production of triketide lactones

Ten mL of each sample was mixed with 2 mL of diethyl ether in a 15 mL conical tube and vortexed for 5 minutes. Each conical tube was centrifuged at 5000g for 10 minutes and 1 mL of ether was removed and placed into a 2 mL flat bottom microcentrifuge tube. Air was gently blown over each sample in a chemical fume hood until dry. The extract was resuspended in 200 uL of methanol with an internal standard of 100 µM δ-nonalactone (Sigma). 5-methyl-6-(propan-2-yl)oxan-2-one was synthesized by Enamine (Cincinnati, USA) to greater than 95% purity used as a standard to approximate 3,5-dimethyl-6-(propan-2-yl)oxan-2-one.

Sample preparation of proteomics

Two milliliters of each production culture were harvested at 10 days, spun down at 10000 g for 10 minutes and resuspended in 0.75 mL of aqueous solution of 50mM potassium phosphate, 300mM NaCl, 10% glycerol and 2 mg/mL of lysozyme at pH 7.5. Samples were incubated for 1 hour at 30 degrees Celsius. Samples were then lysed with the ZR Fungal/Bacterial DNA Microprep kit (Zymo Research, Catalog No: D6007, Irvine, CA). Samples were loaded into the ZR Bashing/Bead Lysis tube and secured in a bead beater. Samples were shaken at a frequency of 30 per second for 5 minutes. The cell lysates were centrifuged at maximum speed in a Eppendorf benchtop centrifuge and the resulting supernatants were collected for proteomic analysis. The protein concentration of protein samples was determined by Bio-Rad DC Protein Assay (Bio-Rad #5000121) according to manufacture instruction. 20 ug protein of each sample was reduced and alkylated, followed by overnight trypsin digestion. The resulting tryptic peptide samples were subjected to LCMS analysis.

LC-MS detection of short chain acids

LC separation of short-chain acids was conducted on an InfinityLab Poroshell HPH-C18 reversed phase column (100 mm length, 3.0 mm internal diameter, 2.7 μm particle size; Agilent, United States) using a Waters Acquity Autopurification system, prep UHPLC-MS (ESI) (Waters, United States). The mobile phase was composed of 10 mM ammonium acetate and 0.05% ammonium hydroxide in water (solvent A) and 10 mM ammonium acetate and 0.05% ammonium hydroxide in methanol (solvent B) to separate 2,4-dimethylpentanoic acid and 2,4-dimethylpent-2-enoic acid. The mobile phase was composed of 0.1% formic acid in water (solvent A) and 0.1%formic acid in methanol (solvent B) to separate 3-hydroxy-2,4-dimethylpentanoic acid and 2,3-dimethyl-3-oxopentanoic acid. All acids were each separated via the following gradient: increased from 5 to 97.1% B in 2.0 min, held at 97.1% B for 2.8 min, decreased from 97.1 to 5% B in 0.4 min, and held at 5% B for an additional 5.8 min. The flow rate was held at 0.42 ml · min−1 for 5.2 min, and then increased from 0.42 to 0.65 ml · min−1 for an additional 5.8 min. The total LC run time was 11 min. Samples were injected into the LC column at a volume of 15 μl. Acids were detected via [M -H]- ions: *m*/*z* = 129.092; *m*/*z* = 127.076; *m*/*z* =145.087; *m*/*z* =143.071. The analysis was performed using an *m*/*z* range of 70 to 500.

LC-MS detection of triketide lactones

LC separation of triketide lactones was conducted on a Kinetex XB-C18 reversed phase column (100 mm length, 3 mm internal diameter, 2.6 μm particle size; Phenomenex, United States) using an Agilent 1200 Rapid Resolution LC system (Agilent Technologies, United States). The mobile phase was composed of water (solvent A) and methanol (solvent B). Lactones were each separated via the following gradient: increased from 30 to 90% B in 3.7 min, held at 94% B for 5.2 min, decreased from 90 to 30% B in 0.33 min, and held at 30% B for an additional 2.0 min. The flow rate was held at 0.42 ml · $min^{-1}$ for 8.67 min, increased from 0.42 to 0.60 ml · $min^{-1}$ in 0.33 min, and held at 0.60 ml · $min^{-1}$ for an additional 2.0 min. The total LC run time was 11.0 min. The column compartment and autosampler temperatures were set to 50°C and 6°C, respectively. Samples were injected into the LC column at a volume of 3 µl. The Agilent 1200 Rapid Resolution LC system was coupled to an Agilent 6210 TOF (Agilent Technologies, United States). Nitrogen gas was used as both the nebulizing and drying gas to facilitate the production of gas-phase ions. The drying and nebulizing gases were set to 10 l · $min^{-1}$ and 25 l · $bin^{-2}$, respectively, and a drying gas temperature of 325°C was used throughout. Atmospheric pressure chemical ionization was conducted in the positive-ion mode with capillary and fragmentor voltages of 3.5 kV and 100 V, respectively. The skimmer, OCT1 RF, and corona needle were set to 50 V, 170 V, and 4 µA, respectively. The vaporizer was set to 350°C. Lactones were detected via [M + H]+ ions: *m*/*z* = 157.1223; *m*/*z* = 171.1379. The analysis was performed using an *m*/*z* range of 70 to 1100. Data acquisition and processing were performed using MassHunter software (Agilent Technologies, United States).

Chemical synthesis

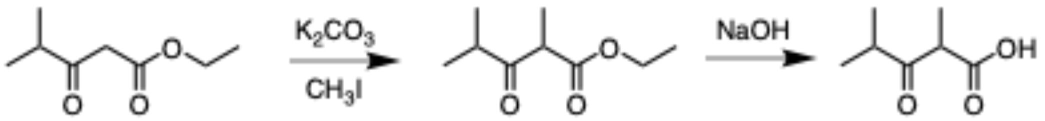


**Synthesis of 2,4-dimethyl-3-oxopentanoic acid**. A solution of ethyl 4-methyl-3-oxopentanoate (1.5 g, 9.5 mmol), potassium carbonate (3.93 g, 28.5 mmol) and methyl iodide (1.62 g, 11.4 mmol) in THF (20 mL) was refluxed overnight under a $N_2$ environment. The reaction mixture was allowed to cool to room temperature before subsequent filtration and concentration was carried out, resulting in the synthesis of ethyl 2,4-dimethyl-3-oxopentanoate as clear yellow oil. Without further purification, the oil was dissolved in 10 mL 1N NaOH aqueous solution and stirred overnight at room temperature. The solution was acidified to pH 1-2 using conc. HCl and the resulting mixture was extracted with diethyl ether (30 mL x 3). The combined organic layers were dried over sodium sulfate, filtered and concentrated *in vacuo* to give the crude product, which was purified by flash column chromatography (silica, DCM: EA 1:1) to yield 2,4-dimethyl-3-oxopentanoic acid a pale-yellow oil (985 mg, 72% over two steps). $^{1}$H NMR (400 MHz, Chloroform-*d*) δ 4.06 (q, *J* = 7.1 Hz, 1H), 2.74 (hept, *J* = 6.9 Hz, 1H), 1.21 (d, *J* = 7.1 Hz, 3H), 1.01 (dt, *J* = 6.9, 2.5 Hz, 6H).

Proteomic analysis

Shotgun proteomic was analyzed on an Agilent 6550 iFunnel Q-TOF mass spectrometer (Agilent Technologies, Santa Clara, CA) coupled to an Agilent 1290 UHPLC system as described previously[32]. Peptides were separated on a Sigma–Aldrich Ascentis Peptides ES-C18 column (2.1 mm × 100 mm, 2.7 μm particle size, operated at 60°C) at a 0.400 mL/min flow rate and eluted with the following gradient: initial condition was 95% solvent A (0.1% formic acid) and 5% solvent B (99.9% acetonitrile, 0.1% formic acid). Solvent B was increased to 35% over 120 min, and then increased to 50% over 5 min, then up to 90% over 1 min, and held for 7 min at a flow rate of 0.6 mL/min, followed by a ramp back down to 5% B over 1 min where it was held for 6 min to re-equilibrate the column to original conditions. Peptides were introduced to the mass spectrometer from the LC by using a Jet Stream source (Agilent Technologies) operating in positive-ion mode (3,500 V). Source parameters employed gas temp (250°C), drying gas (14 L/min), nebulizer (35 psig), sheath gas temp (250°C), sheath gas flow (11 L/min), VCap (3,500 V), fragmentor (180 V), OCT 1 RF Vpp (750 V). The data were acquired with Agilent MassHunter Workstation Software, LC/MS Data Acquisition B.06.01 operating in Auto MS/MS mode whereby the 20 most intense ions (charge states, 2–5) within 300–1,400 m/z mass range above a threshold of 1,500 counts were selected for MS/MS analysis. MS/MS spectra (100–

1,700 m/z) were collected with the quadrupole set to “Medium” resolution and were acquired until 45,000 total counts were collected or for a maximum accumulation time of 333 ms. Former parent ions were excluded for 0.1 min following MS/MS acquisition. The successfully identified peptides of engineered LipPKS1 were targeted in a SRM method developed on an Agilent 6460 QQQ mass spectrometer system coupled with an Agilent 1290 UHPLC system (Agilent Technologies, Santa Clara, CA). Peptides were separated on an Ascentis Express Peptide C18 column [2.7-mm particle size, 160-Å pore size, 5-cm length × 2.1-mm inside diameter (ID), coupled to a 5-mm × 2.1-mm ID guard column with the same particle and pore size, operating at 60°C; Sigma-Aldrich] operating at a flow rate of 0.4 ml/min via the following gradient: initial conditions were 98% solvent A (0.1% formic acid), 2% solvent B (99.9% acetonitrile, 0.1% formic acid). Solvent B was increased to 40% over 11 min, and was then increased to 80% over 1 min, and held for 1.5 min at a flow rate of 0.6 mL/min, followed by a ramp back down to 2% B over 0.5 min where it was held for 1 min to re-equilibrate the column to original conditions. The eluted peptides were ionized via an Agilent Jet Stream ESI source operating in positive ion mode with the following source parameters: gas temperature = 250°C, gas flow = 13 liters/min, nebulizer pressure = 35 psi, sheath gas temperature = 250°C, sheath gas flow = 11 liters/min, capillary voltage = 3500 V, nozzle voltage = 0 V. The data were acquired using Agilent MassHunter version B.08.02. Acquired SRM data were analyzed by Skyline software version 20.1 (MacCoss Lab Software).

Phylogenetic trees

Amino acid sequences of KR and KS domains from 72 BGCs were extracted from ClusterCAD[17]. The sequences were aligned using Muscle v3.8 (https://www.ncbi.nlm.nih.gov/pubmed/15034147)[33]. The alignments were manually curated using JalView (https://academic.oup.com/bioinformatics/article/25/9/1189/203460)[34]. The best amino acid substitution model for both the ketosynthases and ketoreductases phylogenies was

LG+F+I+G4, and it was selected using the ModelFinder tool implemented in IQ-tree[35]. Finally the phylogeny was constructed using IQ-tree[36], using the partitioned models with 10,000 bootstrap replicates and visualized with FigTree (available from http://tree.bio.ed.ac.uk/software/figtree/).

Statistics

Spearman rank correlations were used to compare the significance of reductive loop exchanges. Production titers were normalized to the highest producer in the production of short chain acids in LipPKS1 swaps and triketide lactone production in LipPKS2 swaps. The datasets were combined and production titer and chemosimilarity were normalized to the highest production titer and most chemically similar reductive loop. The datasets were then tested for statistical significance through a Spearman correlation.

Bibliography

1. Phelan, R. M. *et al.* Development of Next Generation Synthetic Biology Tools for Use in Streptomyces venezuelae. *ACS Synth. Biol.* **6,** 159–166 (2017).
2. Yuzawa, S. *et al.* Heterologous Gene Expression of N-Terminally Truncated Variants of LipPks1 Suggests a Functionally Critical Structural Motif in the N-terminus of Modular Polyketide Synthase. *ACS Chem. Biol.* **12,** 2725–2729 (2017).
3. Hagen, A. *et al.* Engineering a polyketide synthase for in vitro production of adipic acid. *ACS Synth. Biol.* **5,** 21–27 (2016).
4. Madeira, F. *et al.* The EMBL-EBI search and sequence analysis tools APIs in 2019. *Nucleic Acids Res.* **47,** W636–W641 (2019).
5. González Fernández-Niño, S. M. *et al.* Standard flow liquid chromatography for shotgun proteomics in bioenergy research. *Front. Bioeng. Biotechnol.* **3,** 44 (2015).
6. Eng, C. H. *et al.* ClusterCAD: a computational platform for type I modular polyketide synthase design. *Nucleic Acids Res.* **46,** D509–D515 (2018).
7. Edgar, R. C. MUSCLE: multiple sequence alignment with high accuracy and high throughput. *Nucleic Acids Res.* **32,** 1792–1797 (2004).
8. Waterhouse, A. M., Procter, J. B., Martin, D. M. A., Clamp, M. & Barton, G. J. Jalview Version 2--a multiple sequence alignment editor and analysis workbench. *Bioinformatics* **25,** 1189–1191 (2009).
9. Kalyaanamoorthy, S., Minh, B. Q., Wong, T. K. F., von Haeseler, A. & Jermiin, L. S. ModelFinder: fast model selection for accurate phylogenetic estimates. *Nat. Methods* **14,** 587–589 (2017).
10. Nguyen, L.-T., Schmidt, H. A., von Haeseler, A. & Minh, B. Q. IQ-TREE: a fast and effective stochastic algorithm for estimating maximum-likelihood phylogenies. *Mol. Biol. Evol.* **32,** 268–274 (2015).
